# Beta-lactam enhancement against methicillin-resistant *Staphylococcus aureus* by cell wall blockade is autolysis-dependent: a butyrolactone derivative as case in point

**DOI:** 10.64898/2026.09.26.754631

**Authors:** Pradnya M. Magdum, Jenny N. Grissom, Brendan T. Franey, Saif Ullah, May H. Abdel Aziz, E. Jeffrey North, Aurijit Sarkar

**Affiliations:** Department of Pharmacy Sciences, School of Pharmacy and Health Professions, Creighton University, Omaha NE U.S.A; Department of Pharmaceutical Sciences and Health Outcomes, Fisch College of Pharmacy, The University of Texas at Tyler, Tyler TX U.S.A

**Keywords:** *Staphylococcus aureus*, MRSA, penicillin, beta-lactam, cell wall, D-Ala-D-Ala, D-cycloserine, autolysis

## Abstract

Methicillin-resistant *Staphylococcus aureus* (MRSA) is non-susceptible to beta-lactams. Blockade of cell wall biosynthesis is a potential target for beta-lactam enhancement but requires further investigation. A butyrolactone derivative enhanced beta-lactams against MRSA strains by reducing the availability of D-Ala-D-Ala. Unlike D-cycloserine, it did not inhibit D-Ala-D-Ala ligase (Ddl). Nor did it show an additive or synergistic effect when combined with cycloserine, indicating a unique mechanism for blocking cell wall precursor production that does not involve the traditional Lipid II pathway. Notably, beta-lactam potentiation by our chemical or D-cycloserine was highly dependent on the intrinsic autolytic ability of the tested MRSA strains. Strains that resisted lysis upon Triton X-100 exposure showed a minimal increase in beta-lactam susceptibility, whereas highly autolytic strains showed significant changes in their beta-lactam MICs. We have thus identified autolytic ability as the Achilles’ Heel in the strategy of targeting cell wall biosynthesis for beta-lactam potentiation.

---

*Staphylococcus aureus* is a pathogen of concern. Beta-lactams covalently bind to penicillin-binding proteins (PBPs), which are responsible for cross-linking the *S. aureus* cell wall. While early beta-lactams were largely effective against staphylococcal infections,^1^ acquisition of the *blaZ* gene encoding beta-lactamase (or penicillinase) by most *S. aureus* strains prevents their use today.^2^ Narrow-spectrum anti-staphylococcal semi-synthetic beta-lactams like oxacillin are effective against penicillinase-producing, methicillin-sensitive *S. aureus* (MSSA). Methicillin-resistant (MRSA) strains emerged when MSSA acquired the *mecA* gene, encoding penicillin-binding protein 2a (PBP2a). PBP2a has a low affinity for most beta-lactams and is able to cross-link the cell wall even when these antibiotics are present; it is the main barrier preventing beta-lactams from functioning.^3^ This has led to increased use of vancomycin and other, newer drugs for therapy, and consequently, clinical cases of resistance have been observed against all these. Yet, beta-lactams offer many advantages, such as a great safety and pharmacokinetic profile,^4^ along with coverage for a multitude of infection sites. Common MRSA strains in the USA have genetic loci that increase their susceptibility to beta-lactams.^5^ Clinical incidences of MRSA infections susceptible to beta-lactams are also known.^2, 6^ These studies point to the encouraging notion that beta-lactams can be reintroduced into the clinic against MRSA, despite penicillinase and PBP2a expression.

Beta-lactam enhancement can occur by blocking (a) beta-lactamase, (b) PBP2a, or (c) cell wall construction. Direct beta-lactamase inhibitors (BLIs, e.g., clavulanate, sulbactam, and tazobactam) are the obvious, clinically proven option for beta-lactam enhancement against MSSA, but no such combinations exist against MRSA.^3, 7^ Suppression of beta-lactamase expression to enhance beta-lactams might be possible,^3, 7, 8^, but this science is still in a nascent stage. PBP2a suppression or inhibition would certainly increase beta-lactam potency against MRSA,^9^ but again, further study is required.

Lipid II biosynthesis, cell wall precursor transport into the extracellular space, or precursor utilization for cell wall construction are promising targets for beta-lactam potentiation.^10, 11^ Limiting the amount of substrate available to PBPs would increase beta-lactam potency. For example, D-Ala-D-Ala ligase (Ddl) inhibitor D-cycloserine and other cell wall synthesis inhibitors synergize with beta-lactams.^12^ Inhibitors of key regulatory signals in the cell wall biosynthetic cascade also enhance beta-lactams.^13^ However, cell wall biosynthesis is a complex process, intimately associated with metabolic pathways. Any enhancers that act in novel ways advance the field. Here, we report the discovery of a chemical that reduces cell wall precursor availability in a unique way.

Chemical ***1*** is a butyrolactonylamino-bis(cyclohexylmethyl-) derivative of alanine that we have named BABCA. Its IUPAC name is (2R)-2-[bis(cyclohexylmethyl)amino]-N-(2-oxooxolan-3-yl)propenamide, **Fig 1**); BABCA and structural analogs ***2***-***20*** (**Fig S1**) were synthesized and chemically characterized (see details in **Supplementary Materials**). Among these, we found that only BABCA was capable of enhancing beta-lactams. We found that BABCA does not alter beta-lactamase expression or *mecA* transcript levels; nor does it affect expression of PBP1-4 (*vide infra*). Instead, BABCA suppresses D-Ala-D-Ala availability without inhibiting Ddl, unlike cycloserine. BABCA showed an autolysis-dependent beta-lactam potentiation, as did cycloserine; strains with a higher predisposition to autolysis also showed greater susceptibility to beta-lactams when BABCA was added. Cycloserine and BABCA do not show additivity. Therefore, BABCA does not affect Ddl, related enzymes along the same pathway, or any of the cell wall cross-linking factors. Instead, it suppresses cell wall synthesis in a novel manner. The exact target of this chemical remains uncertain, but it does suggest that blocking cell wall biosynthesis for beta-lactam enhancement is helpful in *S. aureus* strains capable of autolysis. Further structure-function work will certainly increase its value as a potential beta-lactam enhancer, but it is also key to investigating this new path to beta-lactam enhancement and evaluating its potential. And finally, our results highlight the importance of assessing the autolytic ability of clinical strains to determine the value of cell wall biosynthesis inhibitors for beta-lactam potentiation.

**Fig 1.**
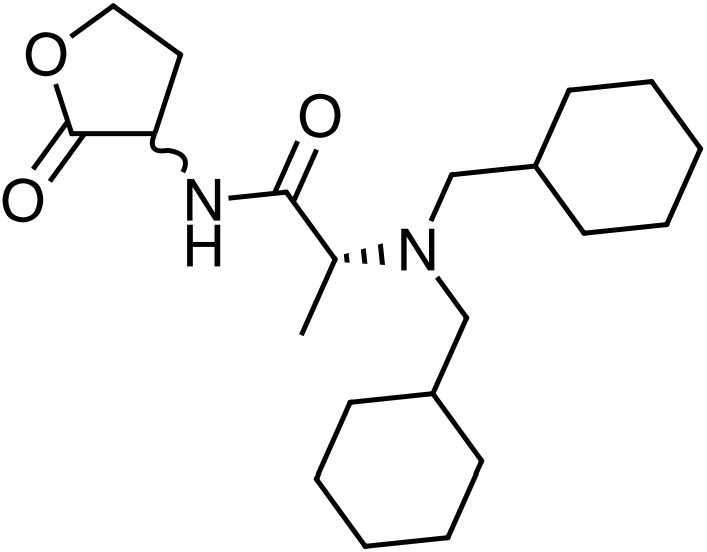
The structure of BABCA. There are two chiral centers, of which one is of undefined stereochemistry. The racemic mixture is tested in this manuscript.

BABCA was synthesized as described in **Scheme 1**, in **Supplementary Methods** and obtained as a racemate. All activities reported are for the racemate. All compounds are 95% pure by HPLC. We studied the effect of BABCA on BAA-1717, a penicillinase-producing USA300 MRSA strain (**Table S1**). BABCA was innocuous to the pathogen by itself (MIC>200 μM, **Table 1**). Higher concentrations of BABCA were not tested because it was dissolved in 100% DMSO, and so we would have >2% DMSO in the vehicle, which could potentially affect bacterial growth. Vancomycin and chloramphenicol were not enhanced by BABCA. Penicillin blocked growth of this strain at 256 μg/mL, as reported in the literature.^8, 13^ 50 μM BABCA increased penicillin potency by 16-fold. Ampicillin and amoxicillin were also enhanced by 8-fold and 4-fold, respectively. Oxacillin, which has a ∼10- to 15-fold reduced affinity for PBP2a when compared with penicillin,^4, 14^ was less enhanced with 50 μM BABCA, but oxacillin was enhanced 8-fold by using 100 μM of the enhancer. So, beta-lactams are enhanced by BABCA to varying degrees, in proportion^3^ to their known affinities for PBP2a and in a concentration-dependent manner. In fact, BABCA is a potent enhancer of beta-lactams; the Clinical & Laboratory Standards Institute M100-S22 document placed the clinical breakpoint of ampicillin-sulbactam combination against penicillinase-producing staphylococci at ≤8 μg/mL of ampicillin in the presence of 4 μg/mL of sulbactam. This is quite close to the ampicillin-BABCA combination in **Table 1**: 32 μg/mL ampicillin stops BAA-1717 growth at 50 μM (∼18 μg/mL) of BABCA.

**Table 1.** MIC enhancement of antibiotics by BABCA and D-cycloserine (D-CS) against BAA-1717.

| Antibiotics | MIC in μg/mL (and fold-change) with different combinations | | | |
| --- | --- | --- | --- | --- |
|  | Control | BABCA<sup>†</sup> | BABCA<sup>@</sup> | D-CS<sup>#</sup> |
| Penicillin | 256 | 16 (16) | 8 (32) | 128 (2) |
| Oxacillin | 256 | 128 (2) | 32 (8) | 16 (16) |
| Ampicillin | 256 | 32 (8) | - | - |
| Amoxicillin | 256 | 64 (4) | - | - |
| Vancomycin | 1 | 1 | 1 | 1 |
| Chloramphenicol | 8 | 8 | 8 | 8 |

#MIC of D-cycloserine against BAA-1717 is 800 μM. Biological replicates (n=2) are presented for all conditions; the more conservative value is reported where needed. Concentrations used: <sup>1</sup>50 μM, <sup>@</sup>100 μM, and <sup>#</sup>200 μM. All controls had a v:v equivalent of DMSO added where appropriate.

^#^MIC of D-cycloserine against BAA-1717 is 800 μM. Biological replicates (n=2) are presented for all conditions; the more conservative value is reported where needed. Concentrations used: ^!^50 μM, ^@^100 μM, and ^#^200 μM. All controls had a v:v equivalent of DMSO added where appropriate.

Cycloserine enhances oxacillin against MRSA.^12^ We investigated whether cycloserine was comparable to BABCA against BAA-1717. Cycloserine showed an MIC of 800 μM (81.67 μg/mL) against this MRSA strain. Here, we will measure cycloserine concentrations in μM rather than μg/mL, since we are interested in it as an enhancer rather than an antimicrobial; this will allow an apples-to-apples comparison with BABCA. As BABCA had an MIC > 200 μM but enhanced beta-lactams at only 50 μM, we checked whether cycloserine has a similar effect. We measured the MIC for varied antibiotics when cycloserine was present at 200 μM, a quarter of its own MIC. Cycloserine enhanced oxacillin by 16-fold, from 256 μg/mL to 16 μg/mL, but it did not enhance penicillin much (**Table 1**). Vancomycin and chloramphenicol were not enhanced. Cycloserine inhibits Ddl to block D-Ala-D-Ala ligation, which inhibits PBP2a function by reducing substrate availability. This results in oxacillin enhancement. BABCA on the other hand, enhanced both beta-lactams, with a preference for penicillin. So, BABCA is mechanistically different.

The fractional inhibitory concentration index (FICI) was calculated to assess for synergy between penicillin and BABCA against BAA-1717 (see **Supplementary Methods**). The MIC of BABCA was set at 200 μM for this calculation, even though the true value is higher (*vide supra*); this would appropriately provide a more conservative estimate of synergy between BABCA and the antimicrobial. The FICI of penicillin and BABCA was 0.3125; thus, BABCA acts synergistically with penicillin against MRSA.

Time-kill curves were performed to evaluate the activity of BABCA alone, and in combination with penicillin G, against BAA-1717 over time (**Fig 2**). Saline was used as a negative control, and vancomycin was used as a positive control, where it exhibited bactericidal activity at just 2 μg/mL, showing a 100-fold reduction in colony-forming units in ≤8 hours. We tested the chemicals at their MIC in kill curves (**Fig S2**), but there was no reduction in bacterial growth. A reduction in number of colonies was observed in the first 8 hours when treated with 4 × MIC of penicillin (1024 μg/mL). Bacterial growth was unaffected when BABCA alone was added at 400 μM. However, when 50 μM of BABCA was added to just 64 μg/mL of penicillin, the kill curve was comparable to 1024 μg/mL of penicillin or 2 μg/mL of vancomycin. BABCA clearly enhances penicillin.

**Fig 2.**
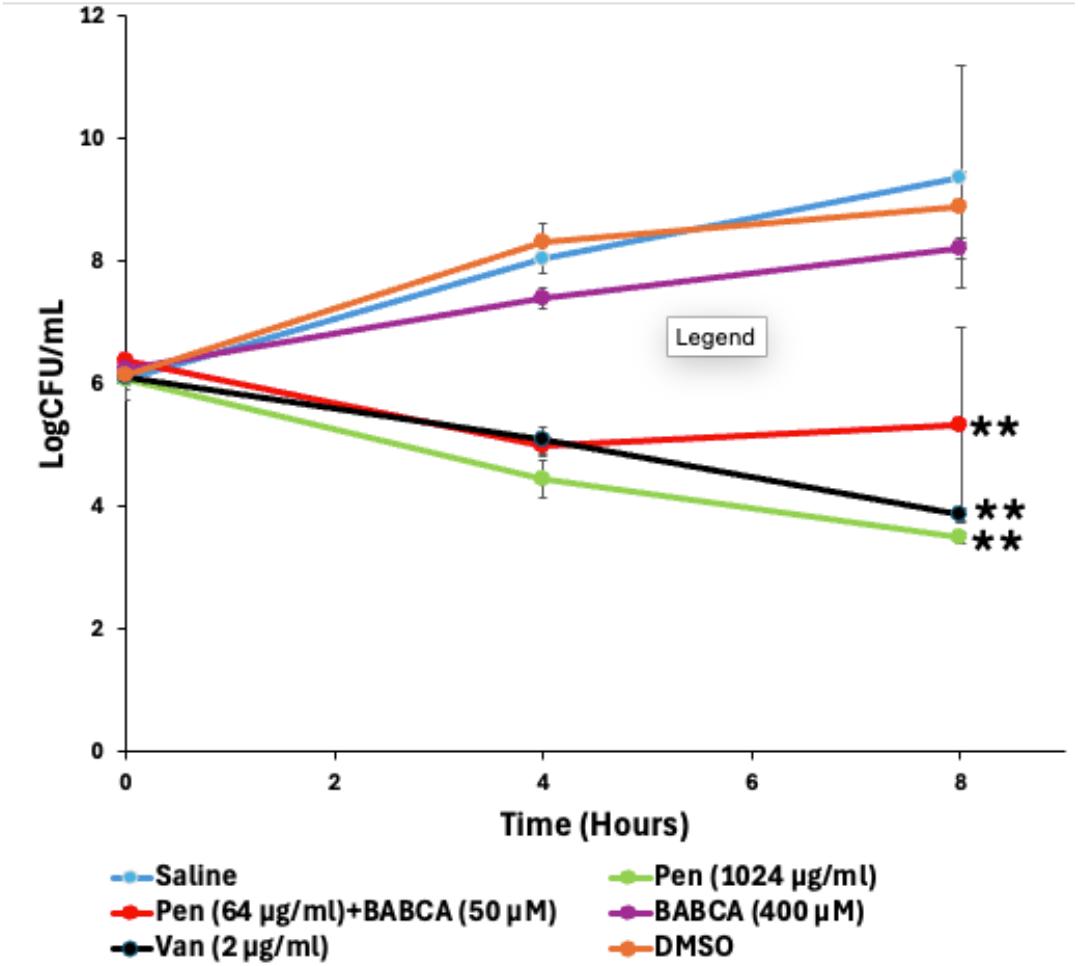
Time-kill curve against BAA 1717. Effect of penicillin G, BABCA and the combination of both compared to the controls. These data represent the mean ± SD from two replicas, and statistical significance was calculated using one-way ANOVA using saline treatment as control (*α = 0*.*01 (**)*).

BABCA has an unusual mode of action. Beta-lactam enhancement may occur because expression of PBPs or PBP2a is reduced or if penicillinase expression decreases, but *blaZ, pbp1-4* and *mecA* gene expression did not change (**Fig 3**, and **Table S2**).^13, 15^ We also evaluated penicillinase activity using a starch-iodide assay^16^ and found no change (**Fig S3**).

**Fig 3.**
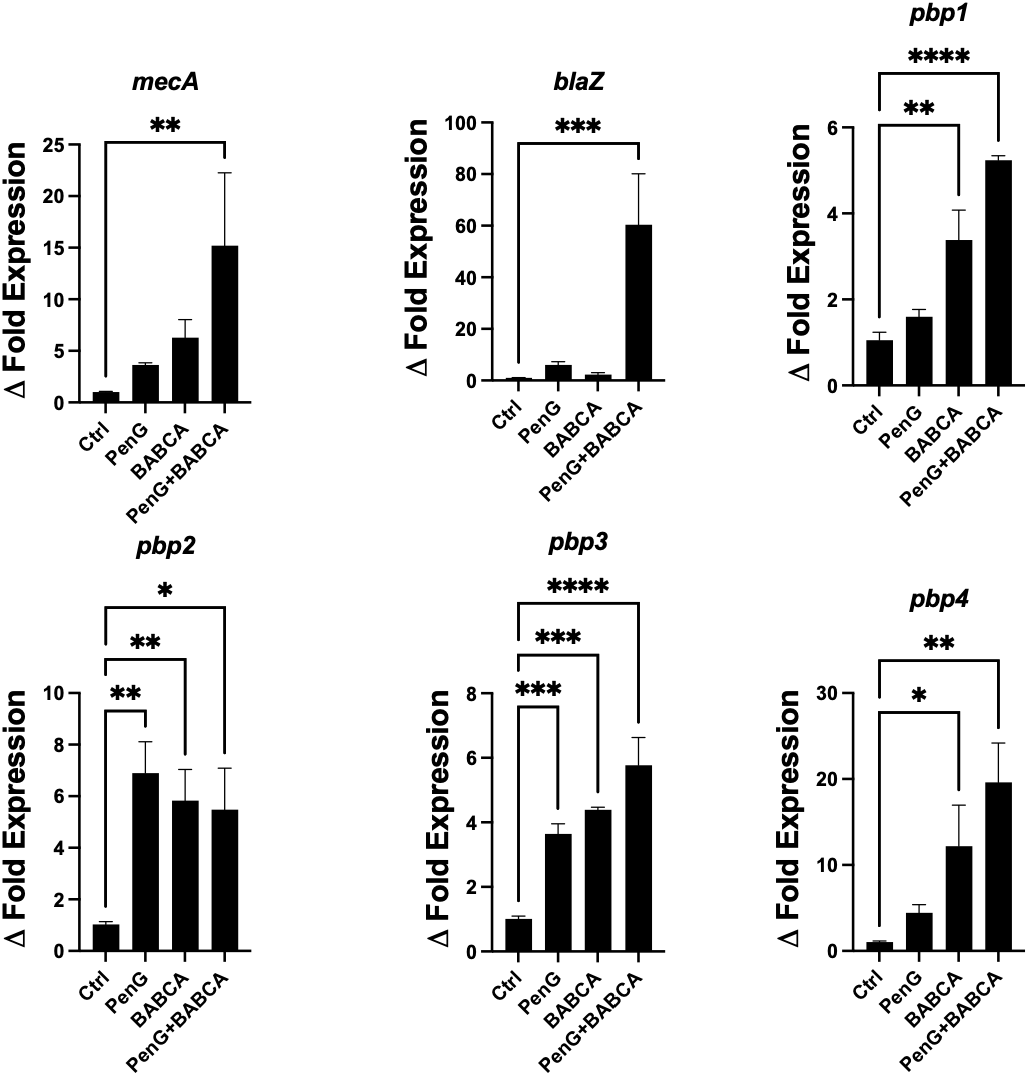
Changes in transcription levels for cell wall biosynthetic and resistance factors in *S. aureus*. The effect of penicillin G (PenG, 128 µg/mL), BABCA (100 µM), and the combination of PenG (8 µg/mL) + BABCA (25 µM) on the fold change of gene expression in the BAA-1717 strain compared to the untreated control (Ctrl). 16S rRNA was used as the reference gene. These data represent the mean ± SEM from at least three replicas, and statistical significance was calculated using one-way ANOVA with multiple comparisons (*α = 0*.*05 (*), 0*.*01 (**), 0*.*001 (***), and 0*.*0001 (****)*).

We investigated whether BABCA exposure might lead to increased autolysis since blockade of autolysis is an inherent defense mechanism against cell wall damaging conditions.^17^ This was not the case for our chemical, however, as **Fig 4** shows there was no increase in autolysis for BAA-1717 when exposed to BABCA or penicillin alone, or a combination of both, BABCA and penicillin. Untreated BAA-1717 cells underwent rapid autolysis over the period of 4 hours when exposed to an autolysis-inducing buffer containing 0.05% Triton X-100 in Tris at pH 7.2 (**Fig 4**, see **Methods** for details); ∼80% reduction in OD580 was observed. As expected, minimal autolysis was observed when untreated cells were exposed to Tris at pH 7.2 (negative control). The difference between these two conditions reflects the *inherent autolytic ability* of this *S. aureus* strain. Importantly, autolysis reduced drastically when BAA-1717 was treated with penicillin, which is a well-known effect.^11^ BABCA, as well as a combination of penicillin and BABCA also reduced autolytic ability.

**Fig 4:**
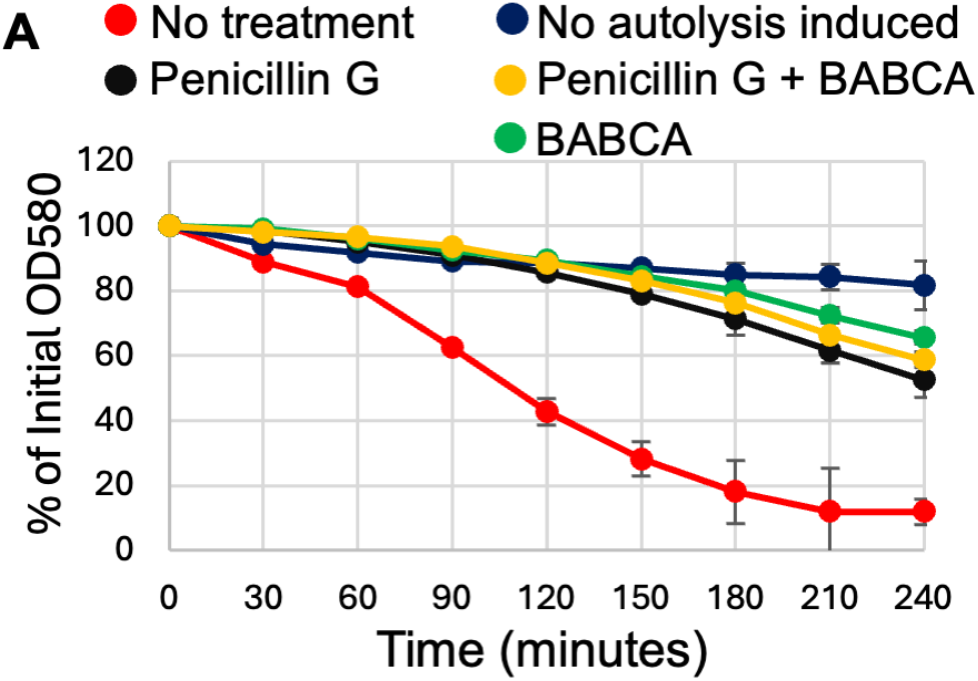
BABCA, penicillin and their combination reduce Triton X-100 induced autolysis in BAA-1717 MRSA. MRSA cells were exposed to various conditions before exposure to autolysis buffer. The “no treatment” sample (red) shows the inherent autolytic ability of the BAA-1717 strain. Statistical significance was ascertained using single factor ANOVA and post-hoc paired two-way t-tests (α=0.05, n=2).

Given that cycloserine and BABCA enhance beta-lactams, and that cycloserine is a Ddl inhibitor that reduces D-Ala-D-Ala availability in *S. aureus*, we next sought to identify whether BABCA works by a similar mechanism. We measured FITC-vancomycin binding to BAA-1717 after incubating the cells with BABCA and controls to assess whether BABCA alters the availability of D-Ala-D-Ala in *S. aureus*. Fluorescence was measured (λ^ex^, 495nm; λ^em^, 515nm) and normalized, where DMSO was set at 100% availability of D-Ala-D-Ala and cycloserine at 0% (**Table 2)**. Treatment with BABCA reduced FITC-vancomycin binding, as determined by fluorescence intensity measurements. Only 32% D-Ala-D-Ala was available after treatment with BABCA. On the other hand, when treated with chloramphenicol, which is not a cell wall-acting antibiotic, the availability of D-Ala-D-Ala was comparable to DMSO. Together, these results indicate that BABCA significantly decreases the availability of the D-Ala-D-Ala motifs in the bacterial cell wall. However, it does not inhibit Ddl, which is also supported by additional data, below.

**Table 2.** BABCA significantly alters the availability of D-Ala-D-Ala in BAA-1717.

| Treatment | % of D-Ala-D-Ala available for binding FITC-vancomycin |
| --- | --- |
| DMSO | 100 |
| Cycloserine (50 μM) | 0 |
| BABCA (50 μM) | 31.7 ± 22.1 |
| Chloramphenicol (8 μg/mL) | 103.8 ± 25.0 |

The data represents n=2, reported as mean ± SD

We identified that BABCA is a poor fit for the cycloserine-binding pocket of Ddl using docking simulations using the *S. aureus* Ddl structure available on the RCSB Protein Data Bank (PDB code 7u9k). This structure is bound to phosphocycloserine, the active metabolite that inhibits Ddl. A water molecule (w563), E16 and E101 formed a strong hydrogen bonding network with a deprotonated 4-amino moiety on phosphocycloserine (**Fig 5A**). These features were required for successful docking. See **Supplementary Materials** for additional details. The structure was prepared for docking simulations as described in **Supplementary Methods**.^18^ We first validated our docking simulations by redocking phosphocycloserine. We proceeded to study how BABCA may bind Ddl using this optimized docking protocol. We constructed the structure of BABCA in two stereoisomeric states to simulate the chiral center whose configuration is unknown. Both stereoisomers were reproducibly docked into the binding site (**Fig 5B-C**) in triplicate docking runs.

**Fig 5:**
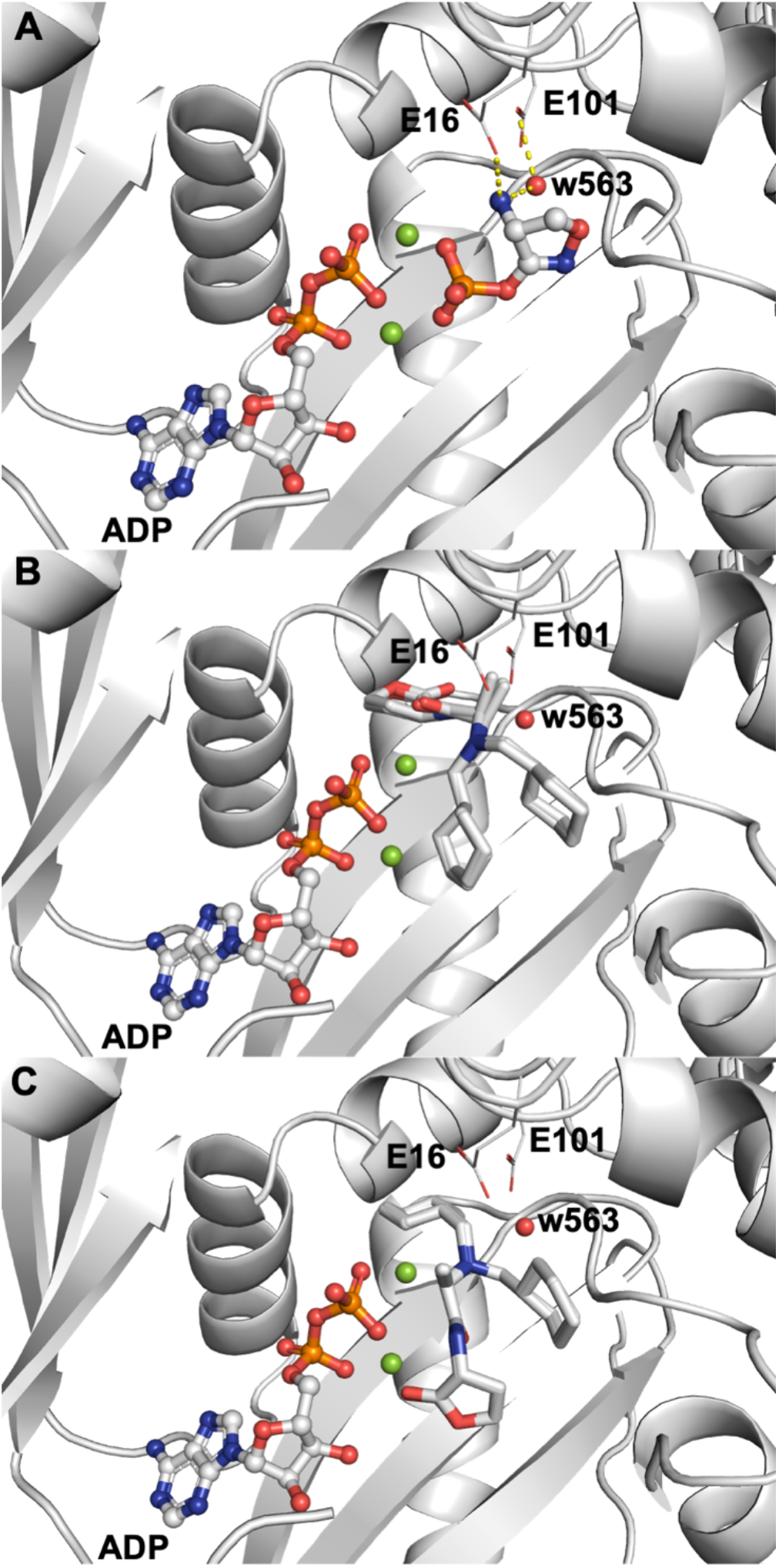
A comparison of phosphocycloserine and docked poses of BABCA within the Ddl orthosteric site. **(A)** Ddl is shown in white ribbons. Phosphocycloserine and ADP are shown in ball-and-stick form (red: oxygen; orange: phosphorus; blue: nitrogen). Mg^2+^ ions are shown as green spheres. A conserved water (w563) is shown as a red sphere. **(B)** and **(C)** show docked poses of two possible configurations of BABCA in stick form with white carbons.

So, we then proceeded to compare the co-crystallized phosphocycloserine ligand’s binding pose with the docked poses of BABCA. While phosphocycloserine presents a primary amine that acts as a hydrogen bond donor to w563, the docked pose of BABCA presents a tertiary amine instead, which makes hydrogen bonding with w563 and E16 difficult. Furthermore, while the phosphate of the co-crystallized ligand interacts favorably with the two Mg^2+^ ions, the two docked poses of BABCA present either the cyclohexylmethyl-substituent to both ions (**Fig 5B**) or the carbonyl oxygen of the butyrolactone substituent to one Mg^2+^ ion and one of the cyclohexyl groups to the other ion (**Fig 5C**). These relatively hydrophobic groups are also in close proximity to the Arg residues nearby, which are also unfavorable interactions. None of the hydrogen bonds from **Fig 5A** are observed with BABCA. Based on these observations, it is evident that BABCA would fail to mimic interactions formed by cycloserine phosphate, which strongly suggests BABCA will not bind Ddl.

We also experimentally supported our predictions from docking. If BABCA inhibits Ddl or a different step in the cell wall biosynthesis pathway, we would expect additive or synergistic effects, as measured by FICI values. So, we tested the MIC of cycloserine in the presence of 50 μM BABCA against strain BAA-1717, and found it was 800 μM, which is exactly the same as with the control shown in **Table S3**. This corresponds to an FICI∼1.5, indicating that the targets of BABCA and cycloserine do not interact. Similarly, BABCA was indifferent to the presence of cycloserine with the penicillinase-negative MRSA strain COL (not shown).

At this point, it was abundantly clear that BABCA does not target traditional targets like Ddl, PBPs or PBP2a. Given this fact, we then asked whether BABCA would function against multiple MRSA strains or if its activity was directed only against BAA-1717, thus reducing its utility. We tested BABCA with beta-lactams against multiple MRSA strains: MW2, N315, COL and Mu50,^17, 19^ which are well-studied, genotyped strains. We found that the MICs of BABCA against MW2 and COL were >200 μM, whereas N315 was 200 μM (**Table S3**), consistent with our observations with BAA-1717 that the chemical does not have significant antimicrobial activity. Mu50 showed growth inhibition by BABCA alone at 100 μM (**Table S3**). This limited the amount of BABCA we could test against these strains for synergy (MW2 and COL, 100 μM; N315, 50 μM; Mu50, 25 μM).

So, we tested for antimicrobial enhancement using 50 μM BABCA against MW2, COL and N315. We found that beta-lactam enhancement by BABCA was entirely strain-dependent (**Table 3**). Penicillin was enhanced ≥4-fold against MW2 and N315 when BABCA was present, but COL did not show increased susceptibility. Even 100 μM BABCA showed only 4-fold enhancement of penicillin against COL. Similarly, 200 μM cycloserine – a known enhancer of oxacillin – enhanced oxacillin only against N315 and MW2, but not against COL and Mu50. Vancomycin and chloramphenicol MICs were not altered by the presence of BABCA in any concentration. No antibiotic was enhanced by BABCA against Mu50 even at ½*MIC (**Table 3**). This shows Mu50 is even less susceptible to BABCA-induced penicillin enhancement than other strains.

**Table 3.**
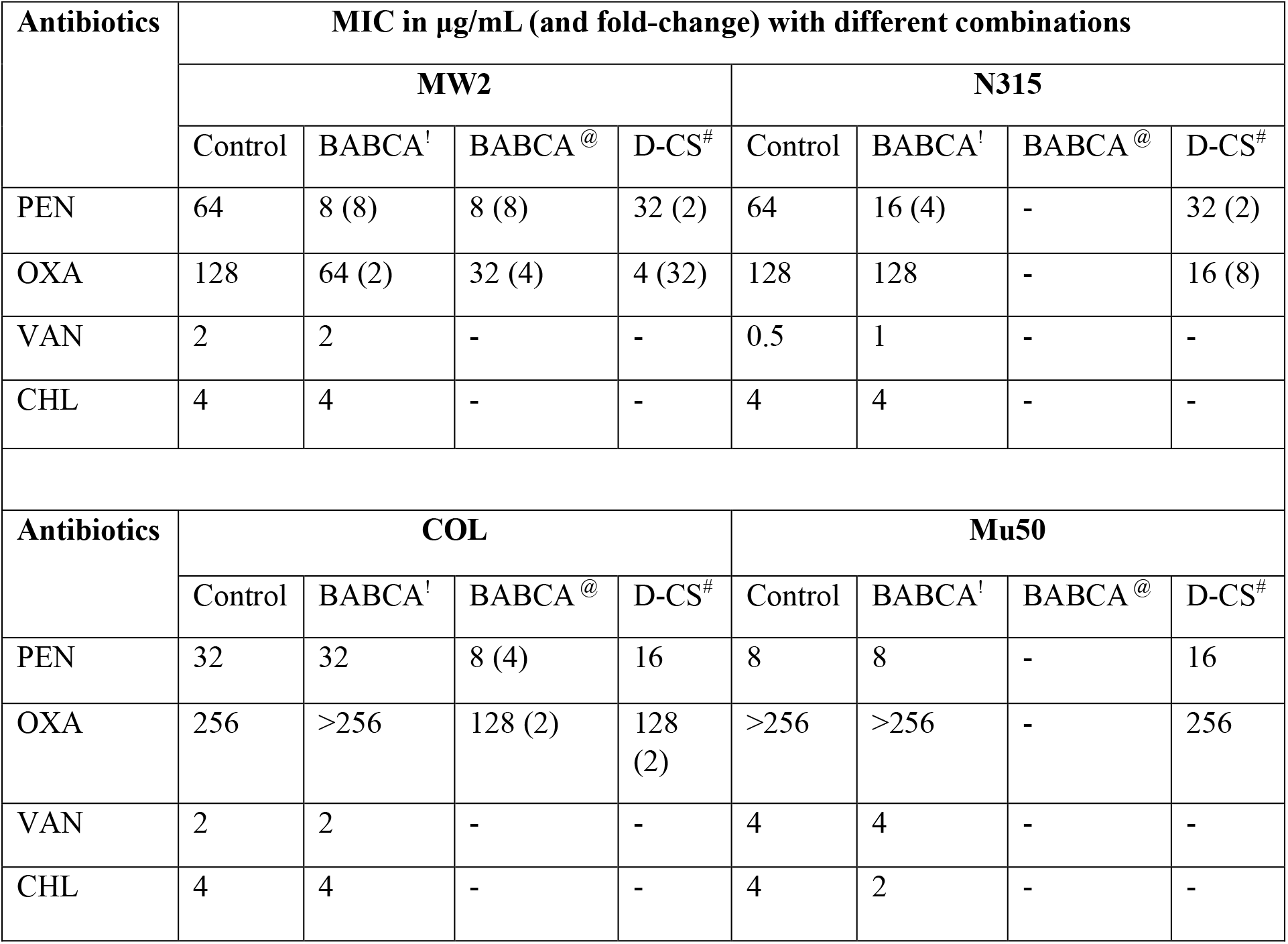
Effect of BABCA and cycloserine on antibiotic susceptibility of tested *S. aureus* strains. Biological replicates were studied (n=2); where replicates showed two separate values, the more conservative MIC is reported to avoid overestimating synergism. Antibiotics: penicillin (PEN), oxacillin (OXA), vancomycin (VAN), chloramphenicol (CHL), and D-cycloserine (D-CS). Concentrations used: ^!^50 μM, ^@^100 μM, and ^#^200 μM.

| Antibiotics | MIC in μg/mL (and fold-change) with different combinations | | | | | | | |
| --- | --- | --- | --- | --- | --- | --- | --- | --- |
|  | MW2 |  |  |  | N315 |  |  |  |
|  | Control | BABCA<sup>!</sup> | BABCA<sup>@</sup> | D-CS<sup>#</sup> | Control | BABCA<sup>!</sup> | BABCA<sup>@</sup> | D-CS<sup>#</sup> |
| PEN | 64 | 8 (8) | 8 (8) | 32 (2) | 64 | 16 (4) | - | 32 (2) |
| OXA | 128 | 64 (2) | 32 (4) | 4 (32) | 128 | 128 | - | 16 (8) |
| VAN | 2 | 2 | - | - | 0.5 | 1 | - | - |
| CHL | 4 | 4 | - | - | 4 | 4 | - | - |
| Antibiotics | COL |  |  |  | Mu50 |  |  |  |
|  | Control | BABCA<sup>!</sup> | BABCA<sup>@</sup> | D-CS<sup>#</sup> | Control | BABCA<sup>!</sup> | BABCA<sup>@</sup> | D-CS<sup>#</sup> |
| PEN | 32 | 32 | 8 (4) | 16 | 8 | 8 | - | 16 |
| OXA | 256 | >256 | 128 (2) | 128 (2) | >256 | >256 | - | 256 |
| VAN | 2 | 2 | - | - | 4 | 4 | - | - |
| CHL | 4 | 4 | - | - | 4 | 2 | - | - |

The complete absence of antimicrobial enhancement against Mu50 was striking. Since Mu50 is renowned for stunted autolysis, we asked whether antimicrobial enhancement will occur only in strains capable of autolysis. We measured the inherent autolytic ability of other *S. aureus* strains (MW2, COL, and N315) as described for BAA-1717 in **Fig 4**. Literature suggests such strains vary in autolytic ability.^17, 20^ The MRSA strains we studied clustered into two distinct groups; N315 and COL showed <25% autolysis upon Triton X-100 exposure, while MW2 resembled BAA-1717 with much higher autolysis. We assessed whether the increased susceptibility to penicillin under exposure to BABCA, as measured by MIC changes (ΔMIC), correlates with their inherent autolytic abilities (without exposure to antibiotic and/or our chemical). **Fig 6A** shows strains that showed a greater ability to undergo autolysis (BAA-1717 and MW2) also showed 8- to 16-fold enhanced penicillin MIC when exposed to 50 μM of BABCA. In comparison, strains with relatively lower autolytic ability (COL and N315) showed a maximum of 4-fold penicillin enhancement. **Fig 6B** shows a similar graph comparing oxacillin enhancement by 100 μM BABCA between the two categories of *S. aureus* strains. We next asked the question whether other beta-lactam enhancers, such as cycloserine, would behave differently. We plotted the MIC fold changes for oxacillin caused by cycloserine across the two categories of *S. aureus* strains and observed a similar trend emerging (**Fig 6C**); strains more capable of autolysis responded strongly to cycloserine as an enhancer, while other strains showed a reduced response. The fact that BABCA and cycloserine, both, augment beta-lactams in an autolysis-dependent manner is critical in our opinion.

**Fig 6:**
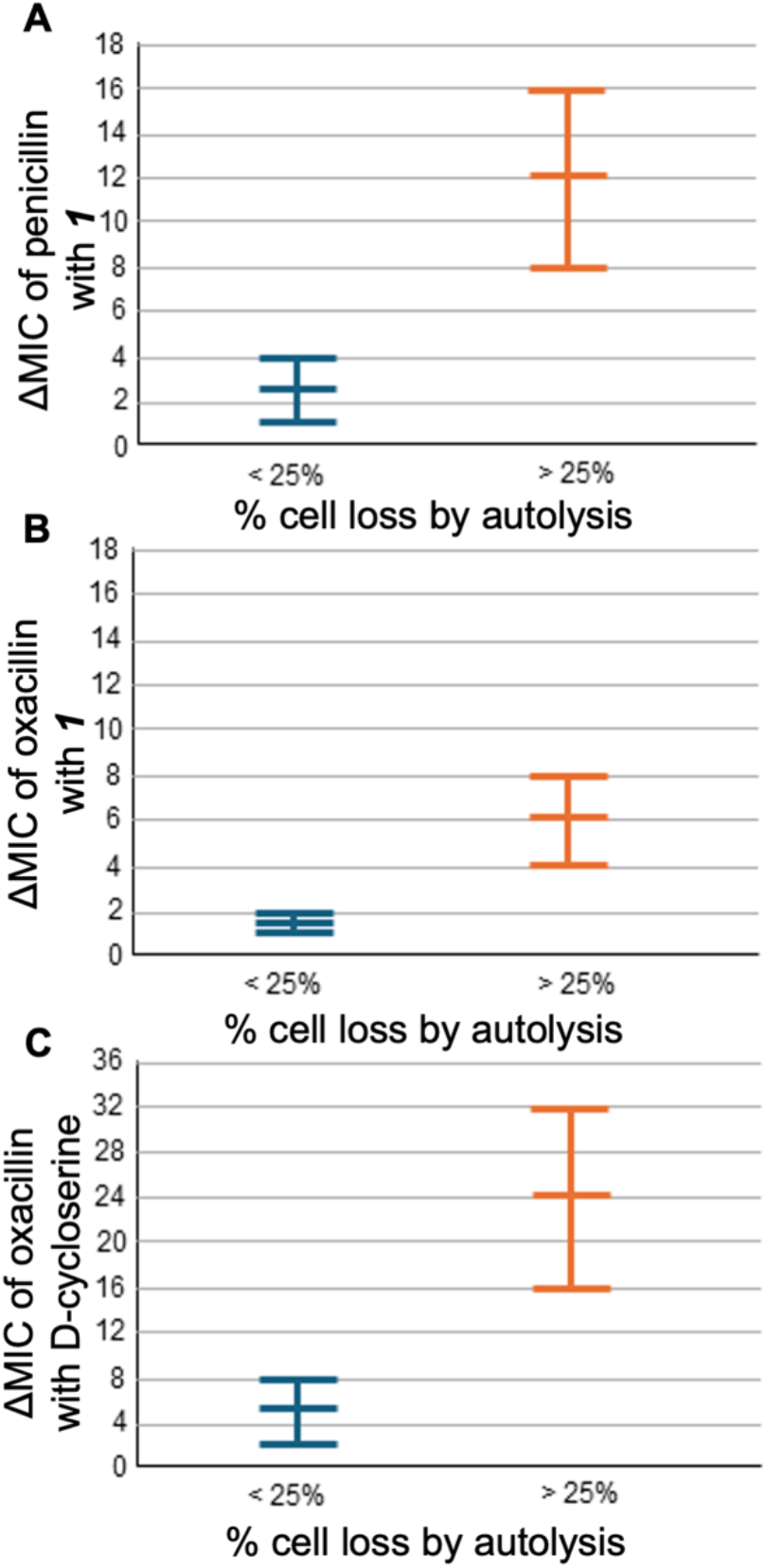
Autolysis determines magnitude of beta-lactam enhancement against *S. aureus* strains. **(A)** MRSA strains with reduced inherent autolytic ability (N315 and COL, with <25% autolytic cell loss upon Triton X-100 exposure) show minimal change in penicillin susceptibility when exposed to 50 μM BABCA. BAA-1717 and MW2 showed >25% autolysis. The complete range of MIC changes is shown. Similar observations were made with the MIC of oxacillin enhanced by 100 μM BABCA **(B)**, or by 200 μM cycloserine **(C)**. A Wilcoxon-Mann-Whitney U test based on n=2 for each condition gave us *p*∼0.06.

## CONCLUSIONS

One of our lab’s major interests is to bring back beta-lactams into the clinic against resistant staphylococci. The rational approach towards achieving this goal is to target resistance factors and regulators. In the past, we have identified a few beta-lactam enhancers that work by suppressing penicillinase expression or by inhibiting regulatory two-component system signaling.^8, 13^ It can also be reasonably expected that blockade of cell wall biosynthesis will synergize with beta-lactams. Here, we present BABCA, a chemical that functions in a unique way.

BABCA is a novel chemical adjuvant capable of restoring the activity of beta-lactams against MRSA. BABCA did not affect the activity of penicillinase or reduce the expression of genes like *blaZ, mecA* or the major PBPs. Instead, it reduced the availability of cell wall precursor D-Ala-D-Ala, suggesting that it perturbs cell wall biosynthesis. And yet, BABCA did not fit into the Ddl binding pocket or show additive effects when coupled with cycloserine; This suggests BABCA does not inhibit the traditional targets for beta-lactam enhancement like Ddl, PBPs or PBP2a. In addition to the potency with which BABCA enhances beta-lactams, the allure of this chemical is also in the indication of a novel (but as yet unknown) pathway that could be exploited to enhance beta-lactams. Moreover, combining such enhancers with more modern drugs like 5^th^ generation cephalosporins, which also have a higher affinity for PBP2a, will also be interesting.

A critical finding was that the beta-lactam enhancement by BABCA (just like cycloserine) depends on the autolytic efficiency of the MRSA strains. Strains with higher autolytic ability exhibited greater BABCA- or cycloserine-induced beta-lactam enhancement, while strains with lower autolytic ability responded poorly. These observations identify autolytic competence as an important determinant of the efficacy of cell wall-targeting beta-lactam enhancers. There was already evidence in literature,^11^ suggesting cell wall biosynthesis blocking antimicrobials showed reliance on autolysis, and that autolysis suppression was a natural response to cell wall-damaging antimicrobials. Our work adds to this observation: Even the strategy of cell wall blockade to enhance beta-lactams will be more effective against cells with higher autolytic capability. To the best of our knowledge, there aren’t any large-scale studies to show the proportion of clinical MRSA that show reduced autolysis – this is perhaps a worthy step to take before pursuing the cell wall blockade strategy.

## AUTHOR INFORMATION

### Author contributions

Conceptualization – PMM, MHA, AS

Methodology – PMM, JNG, BF, EJN, MHA, AS

Data collection – PMM, JNG, BF, SU, AS

Formal analysis – PMM, MHA, AS

Writing and editing – all authors

All authors have read and agreed to the published version of this manuscript.

### Notes

The authors declare no competing financial interests.

## SUPPORTING INFORMATION

^1^H and ^13^C NMR spectra for all chemicals, detailed methods and additional information is provided (PDF).

## ACKNOWLEDGEMENTS

This work was supported by Creighton University start-up funds to AS. SU was supported by the Fisch College of Pharmacy Endowment to MHA. PMM was supported by a Creighton University scholarship. MW2, N315 and COL strains were a gift from Dr. Richard Goering.

