## Supplementary Information for "Beta-lactam enhancement against methicillin-resistant *Staphylococcus aureus* by cell wall blockade is autolysis-dependent: a butyrolactone derivative as case in point"

### SUPPLEMENTARY METHODS

#### *Materials and Reagents*

All the standard antibiotics and chemicals were obtained from Sigma Aldrich, Fisher Scientific and Carolina. We used MRSA and VISA strains (**Table S3**) in this study. BAA-1717 (a.k.a. TCH1516 [USA300-HOU-MR]) is a penicillinase-producing, patient-derived MRSA strain we have studied in the past. MW2, N315 and COL are classical MRSA strains with well-known characteristics. Mu50 is also a well-studied *S. aureus* strain that shows reduced susceptibility to beta-lactams as well as vancomycin. These are key reference strains and their antibiotic susceptibility traits are well-established.

#### *The synthesis of BABCA.*

*Step 1: Synthesis of (1R)-1-[N-2-oxodihydrofuran-3(2H)-ylcarbamoyl]ethyl 2-methylpropane-2-carbamate* (***2***).

Boc-D-alanine (1eq., 1mmol) and DMF (3.0 mL) were stirred at room temperature until dissolved. Under an $N_2$ atmosphere, HOBt (1.5eq., 1.5mmol), EDC (1.5eq., 1.5mmol), and triethylamine (2.5eq., 2.5mmol) were added and left to stir at room temperature for 30 minutes. Dihydro-3-amino-2-(3H)-furanone (1.1eq., 1.1mmol) was added and the $N_2$ atmosphere was recreated and stirred at room temperature overnight. The reaction was concentrated under reduced pressure, and the residue was washed and concentrated with toluene three times. The residue was extracted into DCM and washed with 0.1M NaOH and brine. The organic layer was dried over anhydrous sodium sulfate and evaporated under reduced pressure followed by purification by flash column chromatography.

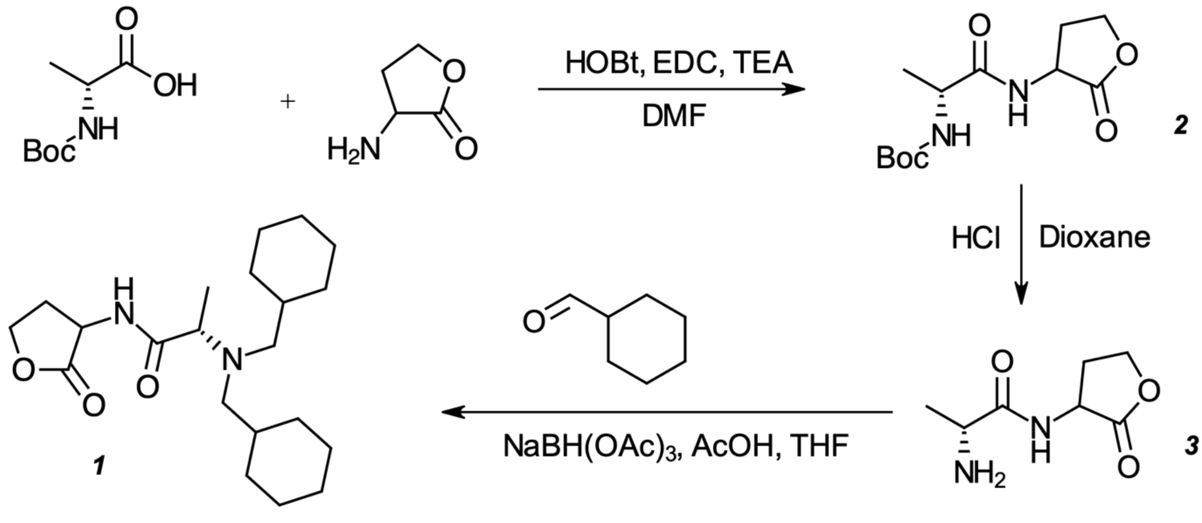


**Scheme 1:** The synthesis of ***BABCA.***

*Step 2: Deprotection of **2** to obtain N-[2-oxodihydrofuran-3(2H)-yl]-(2R)-2-aminopropanamide (**3**).*

***2*** (1eq., 1 mmol) was stirred with 4M HCl in dioxane (4mL) under an $N_2$ atmosphere on an ice bath. After removal from ice bath the reaction was stirred at room temperature and monitored via TLC until completion which took approximately 1 hour. The reaction mixture was then concentrated under reduced pressure to obtain ***3*** and used directly in the next step.

*Step 3: Synthesis of (2R)-2-[bis(cyclohexylmethyl)amino]-N-(2-oxooxolan-3-yl)propenamide* (***BABCA***).

The deprotected substance ***3*** (1.1eq., 0.7259 mmol) was dissolved in THF (3.04 mL) with sonication. Cyclohexanal (1eq., 0.7935 mmol) was added and allowed to stir at room temperature. $NaBH(OAc)_3$ (1.3 eq., 1.03 mmol) was added portion-wise followed by acetic acid and stirred at room temp for 24 hours under $N_2$ atmosphere. The reaction was quenched with saturated sodium bicarbonate and extracted with ethyl acetate and washed with brine. The organic layer was dried over sodium sulfate and concentrated under reduced pressure and the residue was purified with flash column chromatography to obtain BABCA.

*(2R)-2-[bis(cyclohexylmethyl)amino]-N-(2-oxooxolan-3-yl)propenamide* (**BABCA**): 13 mg (8%) of solid; $^{1}$H NMR ($CDCl_3$) δ = 0.63 – 0.93 (4H, m), 1.01 – 1.27 (10H, m), 1.30 – 1.43 (2H, m), 1.49 – 1.74 (8H, m), 1.86 (2H, t, J= 14 Hz), 1.96 - 2.21 (4H, m), 2.65 – 2.77 (1H, m), 3.30 – 3.41 (1H, m), 4.16 – 4.26 (1H, m), 4.36 – 4.51 (2H, m), 7.80 (1H, d, J= 36 Hz); $^{13}$C NMR ($CDCl_3$) δ = 6.68, 6.81, 25.64, 26.03, 26.22, 26.71, 29.70, 30.15, 31.54, 31.75, 32.00, 35.91, 48.74, 48.84, 57.92, 59.34, 65.73, 174.76, 175.27; ESI-MS $[M+H]^+$ calculated for $C_{21}H_{36}N_2O_3$ at 365.1.

*tert-butyl N-[(1R)-1-[(2-oxooxolan-3-yl)carbamoyl]ethyl]carbamate* (**2**): 123 mg (45%) of solid; $^{1}$H NMR ($CDCl_3$) δ = 1.41 (3H, d, J = 7 Hz), 1.45 (9H, s), 2.13 – 2.30 (1H, m), 2.73 – 2.85 (1H, m), 4.18 – 4.35 (2H, m), 4.44 – 4.70 (2H, m) 5.10 (1H, d, J= 6 Hz), 7.00 (1H, d, J= 27 Hz); $^{13}$C NMR ($CDCl_3$) δ = 17.84, 18.05, 28.32, 48.95, 49.13, 49.99, 65.95, 173.42, 174.99, 175.11; ESI-MS $[M+H]^+$ calculated for $C_{12}H_{20}N_2O_5$ at 273.2.

*(2R)-2-amino-N-(2-oxooxolan-3-yl)propanamide* (**3**): 70.2 mg (99%) of solid; $^{1}$H NMR ($CD_3OD$) δ = 1.54 (3H, t, J= 8 Hz), 2.24 – 2.43 (1H, m), 2.52 – 2.66 (1H, m), 3.98 (1H, q, J= 7 Hz), 4.29 – 4.38 (1H, m), 4.48 (1H, t, J= 9 Hz), 4.70 (1H, q, J= 10 Hz); $^{13}$C NMR ($CD_3OD$) δ = 16.00, 28.04, 48.75, 65.84, 169.80, 175.55; ESI-MS $[M+H]^+$ calculated for $C_7H_{12}N_2O_3$ at 173.1.

*tert-butyl N-[(1R)-1-(cyclopentylcarbamoyl)ethyl]carbamate* (**4**): 116 mg (61%) of solid; $^{1}$H NMR ($CDCl_3$) δ = 1.23 – 1.35 (5H, m), 1.38 (9H, s), 1.47 – 1.66 (4H, m), 1.83 – 1.96 (2H, m), 3.95 – 4.17

(2H, m),6.10 (1H, s), 4.96 (1H, s); $^{13}C$ NMR ($CDCl_3$) δ = 23.72, 28.30, 33.04, 51.10, 155.61, 172.05; ESI-MS $[M+H]^+$ calculated for $C_{13}H_{24}N_2O_3$ at 257.2.

*(2R)-2-amino-N-cyclopentylpropanamide* (**5**): 38.1 mg (99%) of solid; $^1H$ NMR ($CD_3OD$) δ = 1.44 – 1.58 (5, m), 1.59 – 1.69 (2H, m), 1.70 – 1.82 (2H, m), 1.90 – 2.02 (2H, m), 3.89 (1H, q, J= 7 Hz), 4.10 – 4.18 (1H, m); $^{13}C$ NMR ($CD_3OD$) δ = 16.42, 23.34, 32.08, 48.85, 51.28, 169.04**;** ESI-MS $[M+H]^+$ calculated for $C_8H_{16}N_2O_1$ at 157.2.

*tert-butyl N-[(1R)-1-(cyclohexylcarbamoyl)ethyl]carbamate* (**6**): 168 mg (89%) of solid; $^1H$ NMR ($CDCl_3$) δ = 1.01 – 1.16 (3H, m), 1.23 – 1.33 (5H, m), 1.38 (9H, s), 1.49 – 1.57 (1H, m), 1.58 – 1.69 (2H, m), 1.76 – 1.86 (2H, m), 3.61 – 3.73 (1H, m), 4.01 (1H, s), 4.95 (1H, s), 5.97 (1H, s); $^{13}C$ NMR ($CDCl_3$) δ = 18.25, 24.68, 25.50, 28.32, 32.94, 48.05, 171.47; ESI-MS $[M+H]^+$ calculated for $C_{14}H_{26}N_2O_3$ at 271.3.

*(2R)-2-amino-N-cyclohexylpropanamide* (**7**): 135.3 mg (99%) of solid; $^1H$ NMR ($CD_3OD$) δ = 1.16 – 1.32 (3H, m), 1.32 – 1.42 (2H, m), 1.49 (3H, t, J= 7 Hz), 1.62 – 1.71 (1H, m), 1.74 – 1.83 (2H, m), 1.84 – 1.95 (2H, m), 3.62 – 3.74 (1H, m), 3.86 (1H, q, J= 7 Hz); $^{13}C$ NMR ($CD_3OD$) δ = 16.40, 24.60, 25.13, 32.16, 32.24, 48.49, 48.85, 168.60 ; ESI-MS $[M+H]^+$ calculated for $C_9H_{18}N_2O_1$ at 171.2.

*tert-butyl N-[(1R)-1-[(3-methoxyphenyl)carbamoyl]ethyl]carbamate* (**8**): 168.1 mg (30%) of solid; $^1H$ NMR ($CD_3OD$) δ = 1.36 – 1.50 (12H, m), 2.73 (3H, s), 4.04 – 4.30 (1H, m), 6.69 (1H, d, J= 8 Hz), 7.08 (1H, d, J= 8 Hz), 7.21 (1H, t, J= 8 Hz), 7.30 (1H, s); $^{13}C$ NMR ($CD_3OD$) δ = 17.02, 27.29, 50.87, 54.33, 79.25, 105.75, 109.50, 112.11, 129.14, 139.40, 160.12; ESI-MS $[M+H]^+$ calculated for $C_{15}H_{22}N_2O_4$ at 295.2.

*(2R)-2-amino-N-(3-methoxyphenyl)propanamide* (**9**): 153.4 mg (99%) of solid; $^1H$ NMR ($CD_3OD$) δ = 1.61 (3H, t, J= 7 Hz), 3.80 (3H, s), 4.07 (1H, q, J= 7 Hz), 6.73 (1H, d, J= 9 Hz), 7.12 (1H, d, J= 8 Hz), 7.25 (1H, t, J= 8 Hz), 7.31 (1H, t, J= 2 Hz); $^{13}C$ NMR ($CD_3OD$) δ = 16.19, 49.51, 54.32, 105.76, 109.75, 111.91, 129.34, 138.88, 160.80, 167.80; ESI-MS $[M+H]^+$ calculated for $C_{10}H_{14}N_2O_2$ at 195.1.

*tert-butyl N-[(1R)-1-[(3-hydroxycyclohexyl)carbamoyl]ethyl]carbamate* (**10**): 75.8 mg (40%) of solid; $^1H$ NMR ($CD_3OD$) δ = 0.98 – 1.28 (5H, m), 1.34 (9H, s), 1.39 – 1.54 (2H, m), 1.58 – 1.83 (3H, m), 1.93 – 2.03 (.5H, m), 1.98 (.5H, m), 3.46 – 3.65 (1H, m), 3.81 – 4.04 (2H, m); $^{13}C$ NMR ($CD_3OD$) δ = 17.20, 18.97, 27.27, 31.17, 32.12, 33.93, 38.35, 44.21, 46.78, 50.22, 65.84, 68.26, 79.19, 156.23, 173.61; ESI-MS $[M+H]^+$ calculated for $C_{14}H_{26}N_2O_4$ at 287.2.

*(2R)-2-amino-N-(3-hydroxycyclohexyl)propanamide* (**11**): 77.1 mg (99%) of solid; $^1H$ NMR ($CD_3OD$) δ = 1.08 – 2.00 (10H, m), 2.06 – 2.39 (1H, m), 3.55 – 3.65 (.25H, m), 3.68 – 3.77 (.25H, m), 3.79 – 3.92 (.25H, m), 3.99 – 4.18 (1H, m), 4.96 – 5.06 (.25H, m), 5.38 (.25H, s); $^{13}C$ NMR ($CD_3OD$) δ = 16.31, 16.42, 18.90, 19.10, 20.92, 21.38, 28.40, 29.90, 30.68, 31.26, 32.00, 34.05, 34.63, 34.70, 36.58, 36.66,

38.23, 40.82, 44.34, 44.55, 46.71, 48.84, 65.78, 68.28, 75.44, 75. 49, 75.96, 76.00, 168.58, 168.72, 1168.87; ESI-MS $[M+H]^+$ calculated for $C_9H_{18}N_2O_2$ at 187.2.

*tert-butyl N-[(1R)-1-[(3-hydroxycyclopentyl)carbamoyl]ethyl]carbamate* (**12**): 45.7 mg (24%) of solid; $^1$H NMR ($CDCl_3$) δ = 1.34 (3H, d, J= 7 Hz), 1.45 (9H, s), 1.60 – 1.71 (2H, m), 6.46 (1H, s), 1.95 – 2.35 (4H, m), 5.16 (1H, s), 4.11 (1H, s), 4.37 – 4.50 (2H, m); $^{13}$C NMR ($CDCl_3$) δ = 18.14, 28.31, 30.99, 33.86, 42.69, 49.33, 72.18, 80.18, 155.72, 172.31; ESI-MS $[M+H]^+$ calculated for $C_{13}H_{24}N_2O_4$ at 273.2.

*(2R)-2-amino-N-(3-hydroxycyclopentyl)propanamide* (**13**): 15 mg (99%) of solid; $^1$H NMR ($CD_3OD$) δ = 1.14 – 1.29 (1H, m), 1.37 (3H, t, J= 6 Hz), 1.45- 1.65 (1H, m), 1.72 – 1.96 (2H, m), 2.02 – 2.23 (2H, m), 3.69 – 3.80 (1H, m), 4.20 – 4.31 (1H, m), 5.31 – 5.41 (1H, m); $^{13}$C NMR ($CD_3OD$) δ = 16.26, 29.65, 30.01, 33.00, 38.33, 41.27, 48.85, 49.31, 71.41, 80.17, 156. 52, 169.08; ESI-MS $[M+H]^+$ calculated for $C_8H_{16}N_2O_2$ at 173.1.

*tert-butyl N-[(1R)-1-[(2-hydroxycyclohexyl)carbamoyl]ethyl]carbamate* (**14**): 87.55 mg (46%) of solid; $^1$H NMR ($CDCl_3$) δ = 1.34 – 1.51 (14H, m), 1.53 – 1.68 (4H, m), 1.69 – 1.81 (1H, m), 2.27 (1H, s), 3.85 – 4.01 (2H, m), 4.11 (1H, s), 5.09 (1H, d, J= 41 Hz), 6.26 – 6.62 (1H, m); $^{13}$C NMR ($CDCl_3$) δ = 18.20, 20.03, 23.59, 24.60, 26.95, 28.31, 31.37, 51.02, 68.68, 80.43, 155.80, 172.48; ESI-MS $[M+H]^+$ calculated for $C_{14}H_{26}N_2O_4$ at 287.2.

*(2R)-2-amino-N-(2-hydroxycyclohexyl)propanamide* (**15**): 185 mg (99%) of solid; $^1$H NMR ($CD_3OD$) δ = 1.19 – 2.09 (11H, m), 3.81 – 4.01 (2H, m), 4.02 – 4.163 (.5H, m), 5.24 – 5.49 (.5H, m); $^{13}$C NMR ($CD_3OD$) δ = 16.32, 16.45, 16.54, 19.51, 19.66, 19.81, 23.28, 23.47, 24.25, 26.29, 26.38, 27.59, 31.06, 31.31, 48.85, 51.57, 67.55, 169.09; ESI-MS $[M+H]^+$ calculated for $C_9H_{18}N_2O_2$ at 187.2.

*tert-butyl N-[(1R)-1-[(3-hydroxyadamantan-1-yl)carbamoyl]ethyl]carbamate* (**16**): 176.9 mg (93%) of solid; $^1$H NMR ($CDCl_3$) δ = 1.33 (3H, d, J= 7 Hz), 1.47 (9H, s), 1.51 – 1.63 (2H, m), 1.67 – 1.75 (4H, m), 1.77 – 2.05 (8H, m), 2.25 – 2.32 (2H, m), 4.05 (1H, s), 5.01 (1H, s), 6.03 (1H, s); $^{13}$C NMR ($CDCl_3$) δ = 28.35, 30.60, 34.85, 40.20, 44.07, 48.97, 54.24, 69.16, 155.67, 171.80; ESI-MS $[M+H]^+$ calculated for $C_{18}H_{30}N_2O_4$ at 339.3.

*(2R)-2-amino-N-(3-hydroxyadamantan-1-yl)propanamide* (**17**): 241.3 mg (99%) of solid; $^1$H NMR ($CD_3OD$) δ = 1.48 (3H, d, J= 7 Hz), 1.54 – 1.66 (2H, m), 1.66 – 1.75 (4H, m), 1.91 – 2.02 (6H, m), 2.22 – 2.29 (2H, m), 3.84 (1H, q, J= 7 Hz); $^{13}$C NMR ($CD_3OD$) δ = 16.66, 30.63, 34.64, 39.60, 43.48, 49.10, 54.16, 68.13, 168.53; ESI-MS $[M+H]^+$ calculated for $C_{13}H_{22}N_2O_2$ at 238.9.

*tert-butyl N-[(2R)-1-(3-hydroxypiperidin-1-yl)-1-oxopropan-2-yl]carbamate* (**18**): 71.95 mg (38%) of solid; $^1$H NMR ($CDCl_3$) δ = 1.12 – 1.39 (4H, m), 1.40 – 1.74 (11H, m), 1.75 – 2.07 (2H, m), 2.30 (1H, s), 3.10 – 3.37 (1H, m), 3.40 – 3.59 (1H, m), 3.63- 4.02 (2H, m), 4.64 (1H, s), 5.52 (1H, d, J= 30 Hz); $^{13}$C NMR ($CDCl_3$) δ = 18.89, 19.39, 21.05, 22.15, 22.56, 23.06, 28.39, 32.03, 42.43, 43.05, 45.59, 45.86,

46.33, 49.06, 52.20, 65.52, 65.88, 66.47, 79.58, 80.12, 171.72, 172.13; ESI-MS $[M+H]^+$ calculated for $C_{13}H_{24}N_2O_4$ at 273.2.

*(2R)-2-amino-1-(3-hydroxypiperidin-1-yl)propan-1-one* (**19**): 93 mg (99%) of solid; $^1H$ NMR ($CD_3OD$) δ = 1.43 – 1.77 (5H, m), 1.78 – 2.05 (2H, m), 3.23 – 3.31 (1H, m), 3.35 – 3.93 (4H, m), 4.36 – 4.50 (1H, m); $^{13}C$ NMR ($CD_3OD$) δ = 15.44, 15.66, 20.74, 21.23, 22.38, 22.90, 31.26, 31.76, 32.03, 42.36, 42.80, 45.18, 51.46, 64.99, 65.21, 168.38; ESI-MS $[M+H]^+$ calculated for $C_8H_{16}N_2O_2$ at 173.1.

*(2R)-2-(dibenzylamino)-N-(2-oxooxolan-3-yl)propanamide* (**20**): 32.3 mg (18%) of solid; $^1H$ NMR ($CD_3OD$) δ = 1.11 – 1.25 (3H, m), 2.04 – 2.26 (1H, m), 2.37 – 2.49 (1H, m), 3.23 – 3.36 (1H, m), 3.41 – 3.52 (2H, m), 3.55 – 3.71 (2H, m), 4.14 – 4.25 (1H, m), 4.34 (1H, t, J= 9 Hz), 4.42 – 4.58 (1H, m), 7.11 – 7.18 (2H, m), 7.22 (4H,t, J= 7 Hz), 7.28 – 7.36 (4H, m); $^{13}C$ NMR ($CD_3OD$) δ = 8.52, 28.08, 28.53, 54.01, 57.35, 65.92, 126.94, 128.13, 128.60, 128.77, 176.02; ESI-MS $[M+H]^+$ calculated for $C_{21}H_{24}N_2O_3$ at 353.1.

***Minimum Inhibitory Concentrations (MIC) determination***

*For MIC of antibiotics:* The MIC of antibiotics was determined as recommended by the CLSI guidelines, in biological duplicates using the broth microdilution method. MRSA strain BAA1717 was streaked on tryptic soy agar plates from the glycerol stocks stored at -80°C and incubated at 37°C. An isolated colony was resuspended in 20 mL of TSB and incubated in a shaking incubator at 37°C with shaking at 120 rpm overnight. The antibiotic stocks were prepared in 4% sterile saline at the following concentrations: Penicillin G (512 μg/mL), Oxacillin (512 μg/mL), Vancomycin (64 μg/mL), and Chloramphenicol (128 μg/mL). The antibiotics stocks were serially diluted in a 96-well plate in 4% sterile saline. The final volume of antibiotics in each well, obtained after serial dilution, was 100 μL. The overnight cultures were diluted in MHB II to achieve a final concentration of $1e^6$ CFU/mL; 100 μL of this culture was added to each well in the plates, so that the final bacterial concentration was $5e^5$ CFU/mL. 4% saline was used as a negative control. The plates were incubated at 37 °C for 16 hours under aerobic conditions, and the results were observed manually after incubation for the lowest concentration that blocked the growth of the pathogen. This concentration was noted as the MIC.

*For MIC of chemicals:* Where MIC of chemicals was to be determined in lieu of antibiotics, the above procedure was followed with a few changes: Chemical stocks were prepared by suspending the powders in 100% DMSO to a final concentration of 10 mM. The chemicals were diluted using 4% saline to a working concentration of 400 μM and then serially diluted. Then each well was inoculated with an equal volume of MRSA (as above), followed by incubation for 16 hours at 37 °C; A v:v equivalent of 100% DMSO was used as a negative control in each well.

***Antibiotic Enhancement Assay***

The assay was performed in biological duplicates. The MIC procedure for antibiotics (*vide supra*) was used but modified to add chemicals at requisite concentrations. A v:v equivalent of 100% DMSO was used as a control. The plates were incubated at 37°C under aerobic conditions and observed visually after 16 hours of incubation.

#### *FICI calculations*

The FICI value was calculated using the formula:

$$\text{FICI} = \frac{\text{MIC of penicillin in combination}}{\text{MIC of penicillin alone}} + \frac{\text{MIC of } \textbf{chemical} \text{ in combination}}{\text{MIC of } \textbf{chemical} \text{ alone}}$$

#### *Time Kill Curve*

The assay was performed as per the CLSI M26-A procedure for the time-kill method. The assay was performed in biological duplicates. The effect of penicillin G and BABCA were evaluated alone and in combination at different concentrations. Saline and DMSO were used as negative controls, and vancomycin was used as a positive control. Penicillin G was tested at 1×MIC (256 µg/mL) and 4×MIC (1024 µg/mL), BABCA at 200 µM and 400 µM, and the combination of penicillin G and BABCA was added at 16 µg/mL + 50 µM, respectively, and 64 µg/mL + 50 µM, respectively. The positive control vancomycin was added at a concentration of 2 µg/mL.

The overnight cultures of BAA 1717 were diluted in MHB II to a concentration of $5e^5$ CFU/mL and a final volume of 1 mL, also containing the antibiotic and/or chemical at the above concentrations. These diluted cultures were incubated at 37 °C in a shaking incubator at 120 rpm for the next 24 hours. 50 µL samples were withdrawn at 0-, 4-, 8- and 24-hour time-points. Several 10-fold dilutions in MHB-II were performed, and 10 µL of each dilution was plated on TSA plates using spreaders. Inoculated plates were incubated at 37 °C for 16 hours before determining the colony counts.

#### *Quantitative real-time PCR (qRT-PCR)*

Beta-lactam enhancement can occur by blockade of genes associated with cell wall construction or resistance. Therefore, it was of interest to see whether the synergy of BABCA with penicillin could be explained by the expression of these genes. We investigated the changes in *mecA*, *pbp1-4* and *blaZ* expression in BAA-1717 when exposed to BABCA or penicillin alone, or a combination of both chemicals. The relative transcription levels were evaluated by comparing the change in gene expression between different treatments and untreated controls; gene expression was normalized to *16S rRNA* as the reference gene (primers listed in **Table S2**). The conditions tested were: (A) penicillin alone (128 µg/mL), (B) BABCA alone (100 µM), and (C) the combination of penicillin (8 µg/mL) + BABCA (25 µM). A reduced concentration for penicillin and BABCA in the combination is required because they synergize to kill the bacterium.

The culture of MRSA strain BAA 1717 was diluted from an overnight culture in TSB at a 1:100 ratio. The cultures were treated with 1/2×MIC of the test chemicals and in biological duplicates: penicillin G was added at 128 µg/mL, BABCA at 100 µM. The combination treatment contained penicillin G at 8 µg/mL and BABCA at 25 µM. These cultures were incubated at 37 °C for five hours with continuous shaking at 120 rpm. After five hours, the cultures were harvested by centrifuging at 5000×g for 10 min at 4 °C. For RNA extraction, the cells were first lysed by adding lysostaphin at 1 mg/mL and incubating at 37 °C for 30 min. The RNA was extracted using the RNeasy Plant Mini Kit from Qiagen, following the manufacturer's instructions, as described. The concentrations of the extracted RNAs was measured using the ND1000 Nanodrop by Thermo Fisher Scientific. cDNA synthesis was completed using the iScript Advanced cDNA synthesis kit from Bio-Rad as per manufacturer's instructions. PCR was carried out on the T-100 Thermal cycler from Bio-Rad. 500 ng of the extracted RNA was added to the reaction mixture, which contained 4 µL of the 5x iScript Advanced reaction mix, 1 µL of the iScript Advanced Reverse Transcriptase, and RNA template, and nuclease-free water to make up the final volume to 20 µL. The PCR reaction protocol included a reverse transcription step at 46 °C for 20 minutes, followed by RT inactivation at 95 °C for 1 minute. The concentrations of the cDNA obtained were measured using Nanodrop. The quantification of gene expression was performed using the CFX Duet Real Time PCR System from Bio-Rad with SsoAdvanced Universal SYBR Green Supermix. The reaction mixture contained 10 µL of the SsoAdvanced Universal SYBR Green Supermix, forward and reverse primers, DNA template with 10ng of cDNA, and nuclease-free water to make up the volume to 20 µL. The reaction was carried out with first, the Polymerase activation and DNA denaturation at 95 °C for 30 seconds. This was followed by 30 seconds of denaturation at 95 °C, annealing, and plate read at 60 °C; this step was repeated for 40 cycles. Then a melt curve analysis from 65-95 °C with 0.5 °C increments every 2-5 seconds. The primers are listed in **Table S2**. *16S rRNA* was used as the reference gene, and the expression of individual genes was normalized to *16S* for each sample. The $2^{-\Delta\Delta Cq}$ method was used to calculate the relative fold change in expression of genes of interest, comparing different conditions to the untreated control.

#### ***Starch-iodide assay***

A starch-iodide assay was performed to determine if BABCA acts by suppressing penicillinase activity in MRSA. Starch-iodide assay is an iodometric method to indirectly measure the activity of penicillinase. This assay was performed as described previously. Agar plates were prepared with TSB, nutrient agar, 0.2% soluble starch, 2 µg/mL penicillin G and either 50 µM loratadine, 50 µM BABCA, or v:v DMSO, and poured into 35 x 10 mm petri dishes. The overnight cultures were grown and 5 µL of each overnight culture was added in each agar plate and incubated for 16 hours at 37 °C. After overnight incubation, each plate was flooded with 600 µL of freshly prepared phosphate-buffered saline (pH 6.4) containing 3 mg/mL of iodine, 15 mg/mL potassium iodide and 50 mg/mL of penicillin G. The plates were then observed for decolorization.

Each plate contains penicillin at 2 µg/mL, which is enough to induce penicillinase expression, in combination with loratadine as positive control, since the antihistaminic reduces penicillinase activity at 50 µM (Cutrona et al. ACS Infectious Diseases 2019), which we have previously confirmed, a v:v equivalent DMSO as negative control, or BABCA. A rapid decolorization was observed in the plates containing DMSO, suggesting penicillinase-driven degradation of penicillin into penicilloic acid continued unabated. On the other hand, loratadine-containing plates were almost uniformly colored blue/brown, confirming blockade of penicillin hydrolysis. The plate containing BABCA showed increased decolorization compared with DMSO – this correlated with the RT-qPCR data on the *blaZ* gene, which showed an increase in *blaZ* transcription levels when *S. aureus* was treated with the combination of antibiotic and BABCA. These results were also reflected with other strains, including BAA-1717 and N315, as shown in **Fig S3**. This further confirmed that BABCA does not reduce penicillinase activity.

***Autolysis assay***

The autolysis assay was performed as described previously in biological duplicates. The overnight cultures of BAA-1717 were diluted in TSB to an optical density at 600 nm (OD600) of 0.1. The treatments were added immediately upon dilution: penicillin G (256 µg/mL), BABCA (50 µM), and a combination of penicillin G (256 µg/mL) and BABCA (50 µM), and saline for untreated samples. The cultures were incubated at 37 °C for the next 3 hours with continuous shaking at 250 rpm. After 3 hours of incubation, the cells were harvested by centrifugation at 4100 rpm for 10 minutes at 4 °C. The cells were washed with cold sterile water twice, and the pellet was resuspended in 50 mM Tris (pH 7.2) with or without 0.05% v/v Triton X-100 to an optical density of 0.8 at 580 nm. The samples were incubated at 37 °C with continuous shaking at 250 rpm and their absorbances were recorded every 30 minutes for the next 4 hours. The results were plotted as % of initial $OD_{580}$. The assay was also performed with other MRSA strains for a comparative study.

#### ***Quantification of D-Ala-Ala***

For this experiment, the overnight cultures were treated with the test and control chemicals: 200 µM cycloserine as a positive control, a v/v equivalent of DMSO and 8 µg/mL of chloramphenicol as negative controls, and 200 µM of ***BABCA.*** The cultures were grown for 16 hours, and the cells were harvested by centrifugation at 4100 rpm for 10 minutes. The cells were resuspended in sterile water and adjusted to an optical density (OD) of 0.8 at 600 nm. In a 96-well plate, the FITC-vancomycin stock solution (1 mg/mL) was added to achieve a final concentration of 16 µg/mL in a total volume of 100 µL. To each well, 100 µL of the resuspended culture was added. The plate was kept at room temperature for the next 30 minutes with continuous agitation. After 30 minutes, the fluorescence intensity of the plate was measured at 495 nm excitation wavelength and 515 nm emission wavelength using the Infinite M Nano + plate reader (Tecan, Grödig, Austria).

#### ***Molecular modeling***

The structure of *S. aureus* Ddl was taken from the RCSB Protein Data Bank (PDB) code *7u9k*, chain B; this chain includes 329 out of 356 Ddl amino acids and also has D-Ala-D-Ala, ATP and two $Mg^{2+}$ ions bound. The structure has a 2 Angstrom resolution. Residues 1-3, 32-35, 68-73, 80-85 and 350-356 were not modeled in the crystal structure, but do not form part of the orthosteric site. This structure was compared with *E. coli* ortholog (PDB code *4c5a*), which has cycloserine phosphate – the active metabolite of cycloserine – bound to the orthosteric side, and a 1.65 Angstrom resolution. While Ddl in *7u9k* has ATP bound to it, the same in *4c5a* has ADP, having transferred its terminal phosphate onto cycloserine. A close inspection of both structures demonstrated only slight movements of amino acid residues with a backbone RMSD of 0.97 Angstrom and an all-atom RMSD of 1.1 Angstrom; these calculations were performed using PyMOL Molecular Graphics System v. 3.1 (Schrödinger, LLC). Any perceived atomic movements between the structures were within experimental error, suggesting the Ddl structure in *7u9k* was appropriate for docking studies. We chose to work with a model comprising the protein from *7u9k* bound to ADP, representing the inhibited conformation of Ddl.

We used Autodock Vina v. 1.1.2 for docking and scoring. This version of Vina uses hydrogen bonding, hydrophobic interactions, and a van der Waal's potential, along with an Iterative Local Search optimizer to find good poses for a ligand at a binding site. The binding site comprising the protein from 7u9k, two $Mg^{2+}$ ions, and ADP was prepared using Open Babel v. 3.1 with the atoms being rigid and hydrogens added as per pH~7.0. Any ligands (BABCA and cycloserine) were prepared using Open Babel to be flexible. All ligands were prepared at pH~7.0 or pH~9.0 to simulate different protonation states of the 4-amino group in cycloserine (see the results section for details). Docking parameters were retained at default.

We first redocked cycloserine within the orthosteric site. Our first redocking simulations for cycloserine in its phosphorylated form were carried out with the protein, ADP and two $Mg^{2+}$ ions. Interestingly, simulations assuming pH~7 failed to reproduce the crystallographic position of cycloserine (not shown). Upon closer examination, we found that a water molecule (w563) was positioned in a manner allowing a water-mediated hydrogen bonding network between D-Ala-D-Ala or cycloserine and E101 on Ddl. We found that this water molecule was critical for redocking successfully (residue HOH 563 in *7u9k* or HOH 2020 in *4c5a*); D-Ala-D-Ala formed a hydrogen bond network with E101 in 7u9k through this water molecule. This water-mediated hydrogen bond network is also conserved between E101 and cycloserine in *4c5a*. We further speculated that the 4-amine in phosphocycloserine may not be protonated when bound to Ddl because burying an ionized amine intuitively involves a desolvation penalty. So, we docked phosphocycloserine with and without a protonated 4-amine into the binding site. Indeed, including w563 in the binding site allowed successful redocking of unprotonated phosphocycloserine, but not the protonated molecule, suggesting this unique environment is critical for binding. Cycloserine successfully redocked into the binding pocket, which now includes this conserved water.

After confirming that the docking parameters were appropriate to redocking cycloserine into the binding site, we proceeded to dock the two possible stereoisomers of BABCA into Ddl.

**SUPPLEMENTARY TABLES**

**Table S1:** *S. aureus* strains studied in this manuscript.

| **Strain** | **Phenotype** | **Source** |
|---|---|---|
| BAA-1717 | MRSA | ATCC |
| MW2 | MRSA | Richard Goering (Creighton) |
| N315 | MRSA | Richard Goering (Creighton) |
| COL | MRSA | Richard Goering (Creighton) |
| Mu50 | VISA | ATCC |

**Table S2.** List of primers used in the RT-qPCR experiments.

| Gene | F_primer (5'-3') | R_primer (5'-3') |
|---|---|---|
| *16S* | *CTGTGCACATCTTGACGGTA* | *TCAGCGTCAGTTACAGAC CA* |
| *blaZ* | *GCTTTAAAAGAACTTATTGAGGCTTCA* | *CCACCGATYTCKTTTATAATTT* |
| *mecA* | *CATTGATCGCAACGTTCAATTT* | *TGGTCTTTCTGCATTCCTGGA* |
| *pbp1* | *AGGTAGCGGTTTTGTGTCC* | *TATCCTTGTCAGTTTTACTGTC* |
| *pbp2* | *CAAGCAACAGATCCTCACCCT* | *AATCGCATGGTTTGTTGCCC* |
| *pbp3* | *GTGGACCAACCTCATCTTTA* | *CGGGAGACCCTTATTATTCT* |
| *pbp4* | *CAAGGCATGTACCACCAATG* | *ACGCGGACTATCCAAGAGAA* |

**Table S3.** MIC of BABCA and D-cycloserine against multiple MRSA strains. The MICs from 2 biological replicates are presented.

| **Strain** | **MIC of BABCA** (μM) | **MIC of D-cycloserine** (μM) |
|---|---|---|
| MW2 | >200 | 800 |
| N315 | >200 | 800 |
| COL | 200 | >800 |
| Mu50 | 100 | 800 |

### SUPPLEMENTARY FIGURES

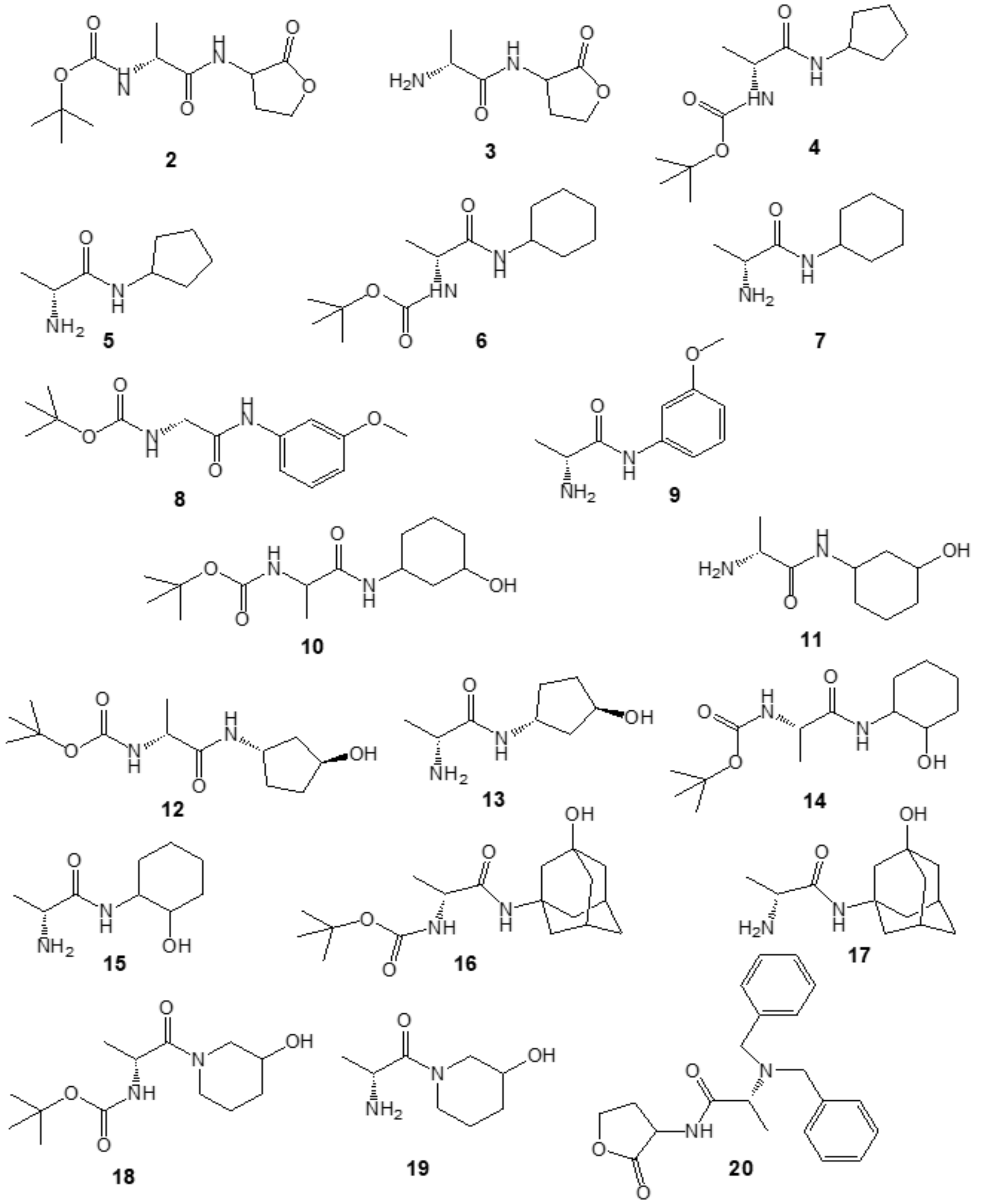


**Figure S1.** Structural analogs of BABCA.

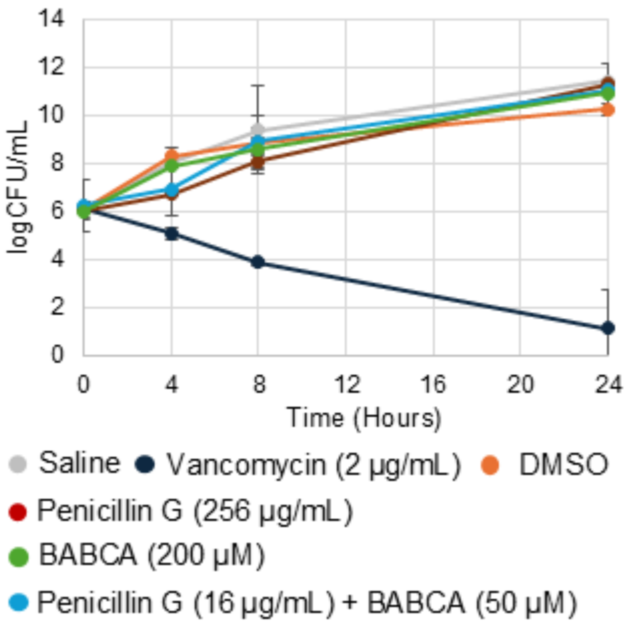


**Figure S2.** Time-kill curves against BAA-1717. The effect of penicillin and BABCA alone and in combination at the concentration of their MICs.

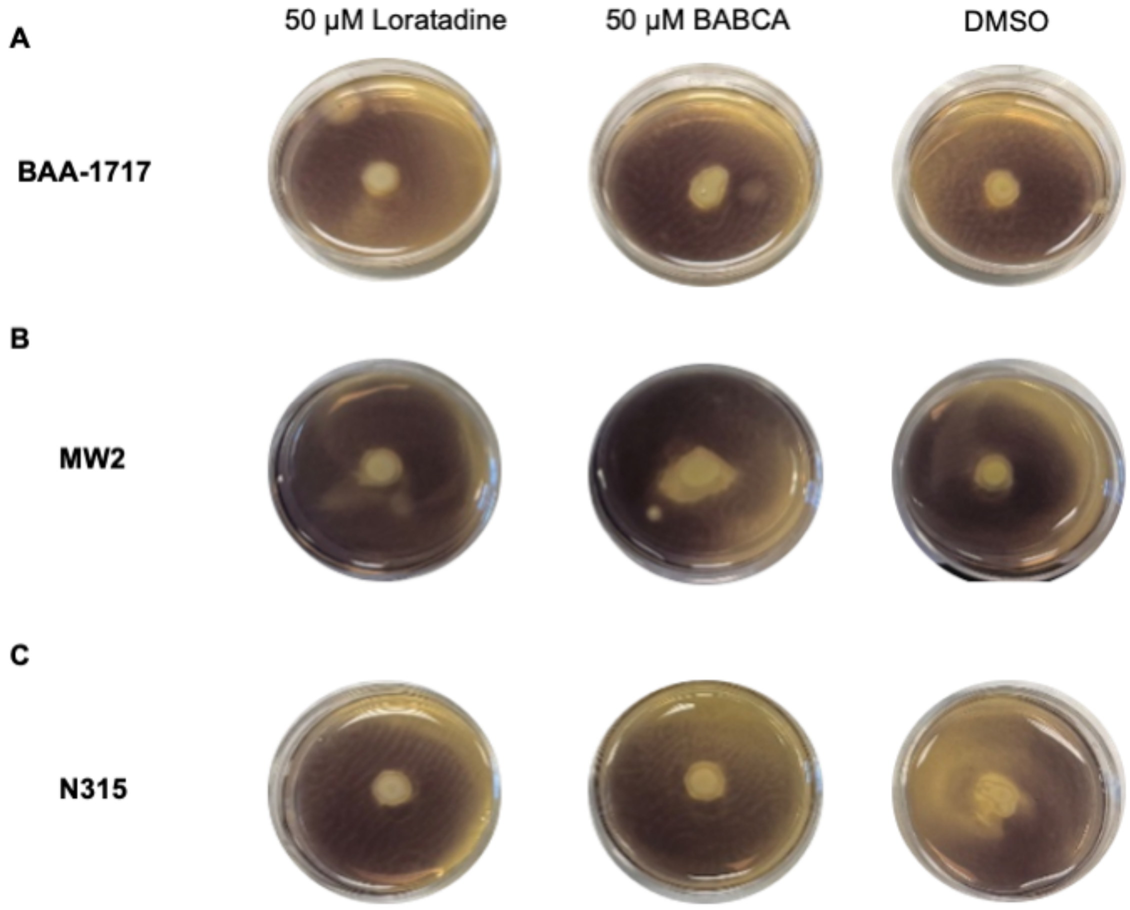


**Figure S3.** Starch-iodide assay to evaluate the effect of BABCA on the expression of penicillinase in strains BAA-1717, MW2 and N315.
